# Differential complex formation via paralogs in the human Sin3 protein interaction network

**DOI:** 10.1101/830828

**Authors:** Mark K. Adams, Charles A.S. Banks, Janet L. Thornton, Mihaela E. Sardiu, Maxime Killer, Cassandra G. Kempf, Laurence Florens, Michael P. Washburn

## Abstract

Despite the continued analysis of HDAC inhibitor efficacy in clinical trials, the heterogeneous nature of the protein complexes they target limits our understanding of the beneficial and off-target effects associated with their application. Among the many HDAC protein complexes found within the cell, Sin3 complexes are conserved from yeast to humans and likely play important roles as regulators of transcriptional activity. The functional attributes of these protein complexes remain poorly characterized in humans. Contributing to the poor definition of Sin3 complex attributes in higher eukaryotes is the presence of two Sin3 scaffolding proteins, SIN3A and SIN3B. Here we show that paralog switching influences the interaction networks of the Sin3 complexes. While SIN3A and SIN3B do have unique interaction network components, we find that SIN3A and SIN3B interact with a common set of proteins. Additionally, our results suggest that SIN3A and SIN3B may possess the capacity to form hetero-oligomeric complexes. While one principal form of SIN3B exists in humans, the analysis of rare SIN3B proteoforms provides insight into the domain organization of SIN3B. Together, these findings shed light on the shared and divergent properties of human Sin3 proteins and highlight the heterogeneous nature of the complexes they organize.

## SUMMARY

Over 13000 or 70% of protein coding genes within the human genome have at least one paralog (1). The acquisition of additional copies of a gene through duplication events provides opportunities for the development of unique gene products with distinct regulatory mechanisms (2). Functional divergence can result from gene duplications and protein paralog identity can influence the composition of large protein complexes (3). However, the consequences of paralog switching are largely overlooked during the characterization of proteins, protein complexes, and protein interaction networks.

Classically associated with transcriptional repression, the removal of histone lysine acetyl groups by the Sin3 histone deacetylase (HDAC) complexes represents a central mechanism whereby transcriptional status is regulated (4). Named for the scaffolding protein of the complex, Sin3 complexes are well studied in *Saccharomyces cerevisiae* (5, 6). However, the presence of additional components not found in lower eukaryotic forms of the Sin3 complexes likely increases the diversity of complex function in higher eukaryotes. Contributing to this expansion of components is the acquisition of paralogous genes encoding Sin3 proteins. The two Sin3 paralogs present within mammals, SIN3A and SIN3B, have undergone significant divergence and maintain only 63% sequence similarity at the protein level in humans.

Genetic deletion of murine *Sin3a* results in early embryonic lethality whereas deletion of *Sin3b* induces late gestational lethality (7, 8). That SIN3A and SIN3B cannot compensate for the loss of one another provides evidence for paralog-specific functions within mammals and suggests that variations of the Sin3 complex have functional consequences. In addition to having critical roles during mammalian development, SIN3A and SIN3B also have contrasting influences on breast cancer cell metastasis. SIN3A was identified as a suppressor of metastasis, whereas SIN3B was proposed to be pro-metastatic (9). However, the mechanisms responsible for divergent influences on development as well as cancer cell metastatic potential remain poorly understood.

SIN3A has been identified as one of the 127 most significantly mutated genes across multiple cancer types (10) and is considered a cancer driver gene (11). Not surprisingly, FDA-approved HDAC inhibitors (HDACis) are effective constituents of multicomponent cancer treatment therapies. While these compounds target the activity of the Sin3 complex catalytic subunits HDAC1/2 (12), heterogeneity within the population of HDAC complexes targeted by these compounds is likely responsible for the well-documented off-target effects associated with HDACis (13) and ultimately prevents our complete understanding of the mechanisms through which these compounds function. The evidence that SIN3A and SIN3B differentially influence metastatic potential, along with the existence of FDA-approved chemotherapeutic agents that target Sin3 complex function, warrants further investigation into the unique properties of complexes containing SIN3A and/or SIN3B as components. Here, we characterize human SIN3A and SIN3B, highlighting the unique and shared properties of Sin3 paralogs in humans. Together, our results shed light on the molecular targets of chemotherapeutic HDAC inhibitors and highlight consequences associated with paralog switching on Sin3 complex composition.

## Results

### HDAC1/2 and RBBP4/7 are core complex components common to SIN3A- and SIN3B-containing complexes

SIN3A and SIN3B share only 63% sequence similarity at the amino acid level (Fig. S1). Therefore, we sought to determine how paralog identity influences protein complex composition. SIN3B isoform 2 (transcript NM_001297595.1, NP_001284524.1) was chosen for analysis as this isoform most closely resembles SIN3A (Fig. S2) and also has strong support as the principal/main isoform within transcriptome databases (https://gtexportal.org/home/gene/SIN3B) (14). To assess differences in SIN3A (NP_001138829.1) and SIN3B isoform 2 interaction networks, open reading frames encoding these proteins were stably expressed in Flp-In-293 cells with C-terminal HaloTags (Fig. 1A, B). Nuclear localization of both recombinant SIN3A and SIN3B isoform 2 (SIN3B_2) was confirmed via fluorescence microscopy (Fig. 1A, B).

**Figure 1.**
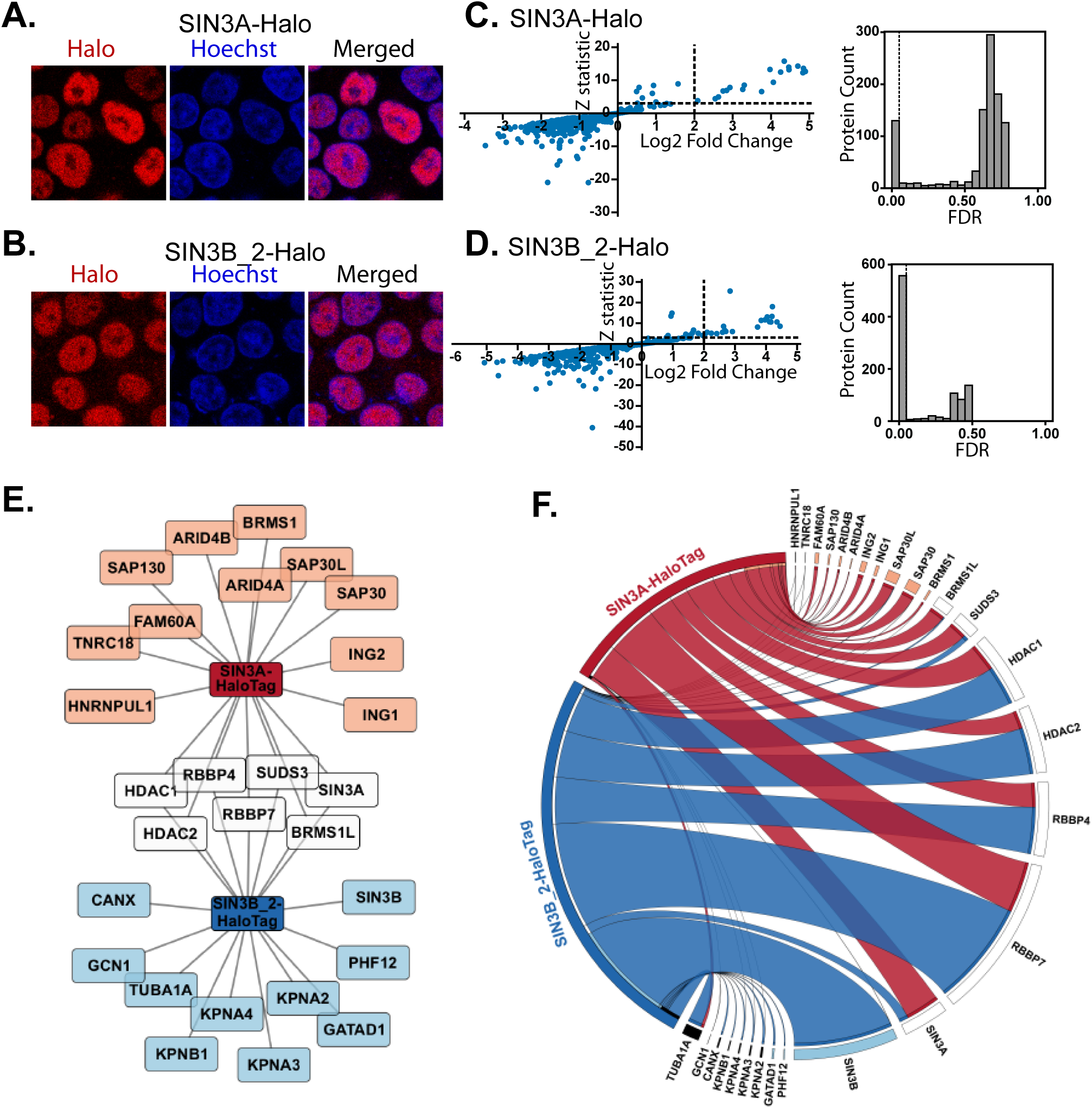
Expression and localization of recombinant Sin3 paralogs and isoforms: (A-B) Subcellular localization of stably expressed (A) SIN3A (NP_001138829.1) and (B) SIN3B_2 (NP_001284524.1) recombinant proteins in Flp-In™-293 cells. HaloTag® TMRDirect™ Ligand and Hoechst 33258 solution were used to visualize recombinant protein localization (red) and nuclei (blue), respectively. (C-D) Significance plots of Z-statistic vs log2 fold change (left panels) and FDR values (right panels) for the proteins detected in each APMS analysis of (C) SIN3A and (D) SIN3B_2 (**Table S2D**). Filter values used for enriched protein identification, Z-statistics ≥ 3 and log2 fold change values ≥ 2, are represented as dashed lines. (E) Network of proteins enriched by SIN3A (SIN3A-HaloTag) and/or SIN3B isoform 2 (SIN3B_2-HaloTag). Recombinant SIN3A and SIN3B are source nodes and proteins enriched by SIN3A (orange), SIN3B_2 (light blue) or both SIN3A and SIN3B_2 (white) purifications are displayed. (F) Circos plot of average dNSAF values for proteins enriched by recombinant SIN3A (red) and/or SIN3B_2 (blue) purifications (Table S2E). For prey proteins with multiple isoforms, dNSAF values for all isoforms were summed and averaged. Prey species meeting enrichment criteria by SIN3A (orange), SIN3B_2 (light blue), or both SIN3A and SIN3B_2 (white) purifications are displayed. With the exception of HDAC1/2 and RBBP4/7, which are known interaction partners of SIN3A/B (55), proteins present in greater than 10% of 411 experiments within the CRAPome database (*Homo sapien* version 1.1) (56) are designated by black bars. *n=3* for SIN3A, *n=4* for SIN3B_2.

Recombinant proteins were affinity-purified and analyzed by MudPIT (Tables S1, S2A-C) as previously described (15). Statistically-significant protein enrichment over negative controls was assessed with QSPEC v 1.3.5 (16) (Fig. 1C,D, Tables S2D-E). Halo-tagged SIN3A captured 18 proteins while SIN3B_2 captured 17 proteins (Fig. 1E, F). While SIN3A and SIN3B_2 enriched a similar number of proteins, the identities of enriched proteins only partially overlapped. Only 7 proteins were enriched in both SIN3A and SIN3B_2 purifications (Fig. 1E).

Within *S. cerevisiae*, 2 forms of the Sin3 complex exist, Rpd3L (Sin3L) and Rpd3S (Sin3S) (5, 6). These complexes share a core of proteins consisting of Sin3, Rpd3, and Ume1, which have homology to human SIN3A/B, HDAC1/2, and RBBP4/7, respectively (17, 18). Using dNSAF values as indicators of protein abundance, HDAC1/2 and RBBP4/7 were identified among the most abundant non-bait proteins in both SIN3A and SIN3B_2 enrichments (Fig. 1F, Table S2F). Thus, SIN3A and SIN3B_2 interact with a set of proteins that are homologous to shared core complex components present within both forms of yeast Sin3 complexes. Interestingly, SIN3A was enriched by both SIN3A and SIN3B_2 (Fig. 1E, Tables S2D-E), suggesting that hetero-oligomeric forms of the Sin3 complexes may exist.

### Both Sin3 paralogs enrich homologs of yeast Sin3L-specific components but only SIN3B enriches a Sin3S-specific component

While sharing a core of subunits, the two yeast Sin3 complexes are differentiated by the identities of additional subunits. Components specifically found within the Rpd3L complex include Sds3, Sap30, and Pho23. These proteins have homology to human SUDS3/BRMS1/BRMS1L, SAP30/SAP30L, and ING1/2, respectively (19–21). Notably, all of these proteins were enriched by SIN3A (Fig. 1E-F). Among these proteins, only SUDS3 and BRMS1L met statistical criteria for enrichment by SIN3B_2 (Fig. 1E). While peptides mapping to BRMS1, ING1/2, and SAP30/SAP30L were observed following SIN3B purification (Fig. 1F), these proteins did not meet criteria for enrichment. These data suggest that SIN3B_2 may interact with at least a subset of proteins with homology to Rpd3L-specific components.

The yeast Rpd3S complex also contains components with homology to human proteins: yeast Rco1 and Eaf3 have homology to human PHF12 and MORF4L1, respectively (22, 23). PHF12 was specifically enriched by SIN3B_2 but not SIN3A. GATAD1, a PHF12 interaction partner was also specifically enriched by SIN3B_2. SIN3A-purified samples were devoid of peptides that mapped to PHF12 (Fig. 1F). Together these data suggest that SIN3A and SIN3B_2 interact with proteins that are homologous to yeast Rpd3L-specific components but only SIN3B_2 interacts with proteins that are homologous to Rpd3S-specific components.

### SIN3A and SIN3B share a common HDAC Interaction Domain (HID)

While isoform 2 likely represents the dominant isoform of SIN3B, the National Center of Biotechnology Information human protein database (June 2016 release) contains 2 additional annotated isoforms. SIN3B isoform 1 (SIN3B_1, NM_015260.3, NP_056075.1) represents the longest isoform and contains an exon absent within isoform 2 (Fig. 2A, S2). Isoform 3 (SIN3B_3, NM_001297597.1, NP_001284526.1) results from an alternative start codon and lacks the N-terminal regions found within isoforms 1 and 2 (Fig. 2A, S2). Since predicted SIN3B protein domains are either absent or altered within these rare SIN3B isoforms (Fig. 2A), we utilized these SIN3B proteoforms to characterize domain organization within SIN3B.

**Figure 2:**
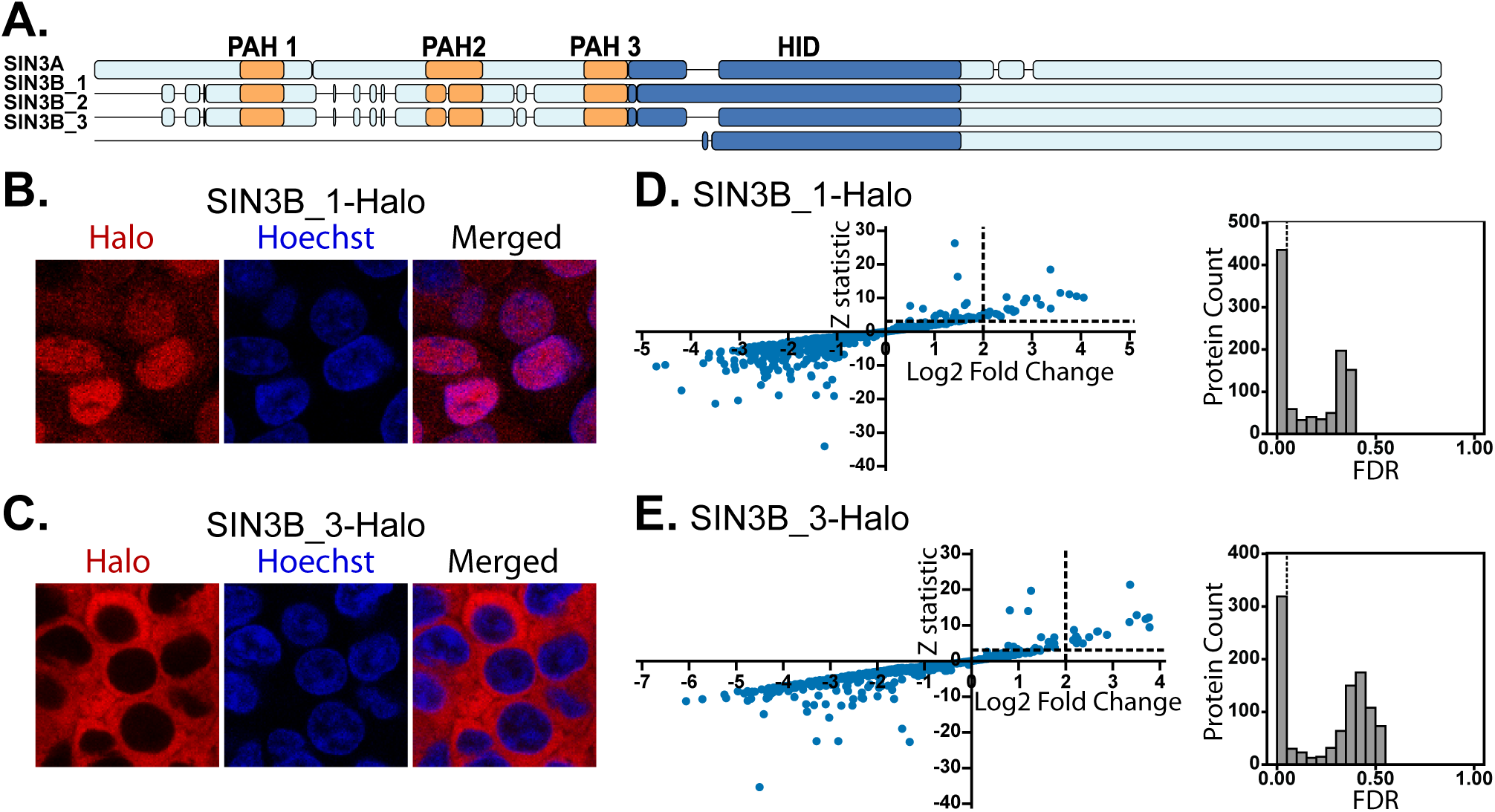
Expression of rare SIN3B proteoforms. (A) Alignment visualization of human SIN3A (NP_001138829.1), SIN3B_1 (NP_056075.1), SIN3B_2 (NP_001284524.1), and SIN3B_3 (NP_001284526.1). PAH domains as defined by Pfam (57) are in orange and the experimentally defined mouse SIN3A HID (24) and homologous sequences in SIN3B isoforms are in dark blue. Detailed sequence alignments are provided in **Figure S2**. (B-C) Subcellular localization of stably expressed (B) SIN3B_1 and (C) SIN3B_3 recombinant proteins in Flp-In-293 cells. HaloTag® TMRDirect™ Ligand and Hoechst 33258 solution were used to visualize recombinant protein localization (red) and nuclei (blue), respectively. (D-E) Significance plots of Z-statistic vs log2 fold change (left panels) and FDR values (right panels) for the proteins detected in each APMS analysis of Sin3 paralogs and isoforms (**Table S2D**). Filter values used to identify enriched proteins are designated by dotted lines.

To assess the characteristics of each isoform, we expressed SIN3B_1 and SIN3B_3 fused with HaloTags in Flp-In-293 cells. Analysis of recombinant protein localization patterns revealed that SIN3B_1 localized within the nucleus (Fig. 2B) whereas SIN3B_3 resided within the cytoplasm (Fig. 2C). Statistical significance of protein enrichment was calculated using QSPEC 1.3.5 (Fig. 2D-E). Hierarchical clustering of protein dNSAF values for proteins enriched by at least one of the four bait purifications revealed that interaction networks of the four Sin3 forms only partially overlapped (Fig. 3A). A group of four proteins was enriched by all analyzed Sin3 proteins (Fig. 3B), consisting of HDAC1, RBBP4, and 2 isoforms of RBBP7.

**Figure 3.**
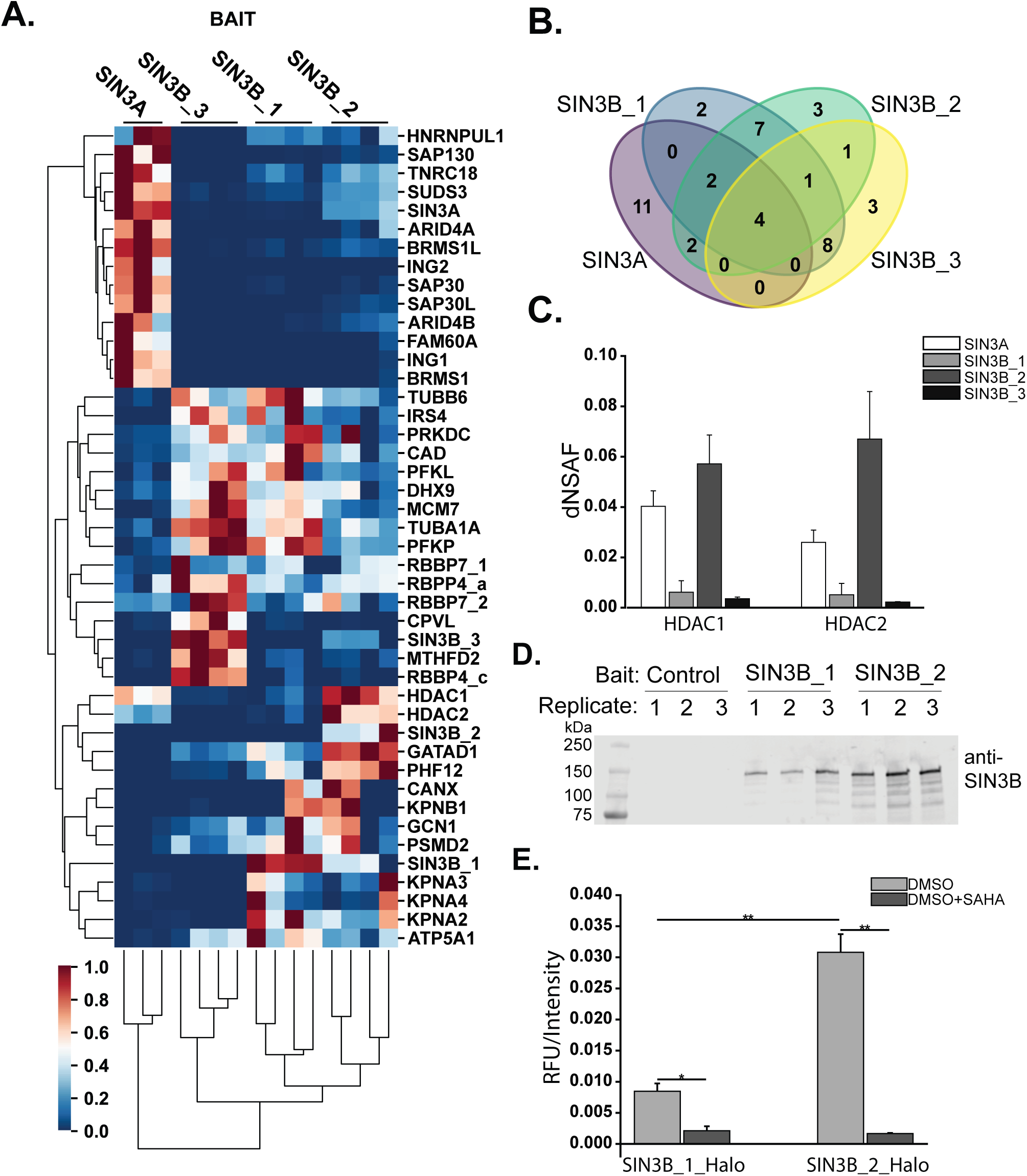
The SIN3A HID is conserved in SIN3B. (A) Clustered heatmap of normalized dNSAF values for proteins within each bait replicate that were enriched by at least one Sin3 protein bait. Proteins were isolated and analyzed from three (SIN3A) or four (all SIN3B isoforms) independent cell clones stably expressing the recombinant Sin3 protein of interest (**Table S2A-E**). Proteins for which multiple isoforms are represented are denoted by isoform identifiers after the protein name. (B) Venn diagram of the number of prey proteins identified as interaction partners of human SIN3A (purple), SIN3B_1 (blue), SIN3B_2 (green), and SIN3B_3 (yellow). (C) Average dNSAF values measured for HDAC1 and HDAC2 in SIN3A-(white), SIN3B_1 (light grey), SIN3B_2 (dark grey), and SIN3B_3 (black) affinity-purified samples. *Mean ± S.D., n=3* for SIN3A, and *n=4* biological replicates for all SIN3B isoforms (**Table S2B**). (D-E) Halo-control (Control), SIN3B_1-HaloTag (SIN3B_1), SIN3B_2-HaloTag (SIN3B_2) were transiently expressed in 293T cells and purified using Magne® HaloTag® Beads. Purified protein was eluted in 100 µL of elution buffer. (D) 10 µL of whole cell extract from each transfection replicate was loaded onto 4-15% polyacrylamide gels and blots were probed using a 1:3000 dilution of anti-mSIN3B sc-13145 X antibody (Santa Cruz Biotechnology, Dallas, TX) and a 1:10,000 dilution of IRDye^®^ 800CW secondary antibody (LI-COR, Lincoln, NE). (E) HDAC activity assay of protein complexes purified using SIN3B isoform 1 (SIN3B_1_Halo) and SIN3B isoform 2 (SIN3B_2_Halo) transiently expressed as baits within 293T cells. Reactions were supplemented with DMSO (grey) or DMSO + SAHA (black). RFU (relative fluorescence unit) values for all biological replicates were normalized to recombinant SIN3B protein abundance in purified samples as measured by Western blot (**Fig. 3D, Table S3**). *Mean ± S.D., n=3*. *: p≤0.005, **: p≤0.001.

Previous studies of mouse SIN3A identified a 327 residue HDAC interaction domain (HID) that is essential and sufficient for interactions with HDAC2 (24). Though a HID within SIN3B has not been experimentally defined, alignment of SIN3A and SIN3B isoforms proteins reveals that a region of SIN3B_2 has high homology to the experimentally defined SIN3A HID (Fig. 2A, S2). The additional exon found within SIN3B_1 resides in the region that aligns with SIN3A HID whereas the HID region in SIN3B_3 is shorter, missing about 1/5^th^ of its N-terminus (Fig. 2A, S2). As this region is highly homologous to the SIN3A HID, we examined what influence the additional exon in SIN3B_1 or the shortened HID in SIN3B_3 had on interactions with the catalytic HDAC subunits of the complex. While HDAC1/2 were enriched by SIN3B_1 and SIN3B_2 (Fig 3A, Table S2E), distributed spectral counts and dNSAF values of these proteins were consistently lower following SIN3B_1 purification (Fig. 3C, Table S2E-F). HDAC1 met criteria for enrichment following SIN3B_3 purification; however, HDAC1 and HDAC2 were less abundant compared to SIN3B_2-purified samples (Fig. 3C, Table S2E-F). Thus, disruption of the HID by the presence of an additional exon in SIN3B_1 or due to shortening of SIN3B_3 N-terminus interferes with, but does not completely inhibit, HDAC1/2 binding.

We next sought to investigate what affect the additional exon present within SIN3B_1 had on the catalytic properties of SIN3B complexes. SIN3B_1 and SIN3B_2 with C-terminal HaloTags were transiently expressed in 293T cells for subsequent protein isolation and HDAC activity assays (Fig. 3D). Activity of the purified protein complexes was assessed with a fluorometric HDAC activity assay. As SIN3B_1 protein levels were consistently lower than that of SIN3B_2 (Fig. 3D), activity was normalized to bait protein abundance (Fig. 3E, S3, Table S3). The enzymatic activity of SIN3B_1-purified samples was consistently lower than purified complexes containing recombinant SIN3B_ 2 (Fig. 3E). Of note, the HDAC activity of both SIN3B complexes was almost completely inhibited by suberoylanilide hydroxamic acid (SAHA) (Fig. 3E). These data suggest that, like the HID region in SIN3A, this region is influences the construction of catalytically active SIN3B-containing complexes.

### The Sin3 HID influences complex assembly

While the HID of Sin3 proteins influences interactions between HDAC1/2 and Sin3 proteins, we next sought to determine what influence alterations to this region in SIN3B isoforms had on other protein-protein interactions.

In addition to HDAC1/2, SIN3A and SIN3B both interact with RBBP4/7. Unlike interactions with HDAC1/2, SIN3B_1 and SIN3B_3 did not display a decreased capacity to interact with RBBP4/7 compared to SIN3B_2 (Fig. 4A, Table 2E-F). The enrichment of RBBP4/7 by SIN3B_3 suggests that the C-terminal half of SIN3B is sufficient for association with these proteins. Of note, RBBP4 isoform c (RBBP4_c, NP_001128728.1) was consistently observed as an interaction partner of SIN3B_ 3 but not of other SIN3B isoforms (Fig. 4A, Table S2E-F). cNLS mapper predicts that a nuclear localization signal with a score of 3.8 is present in RBBP4 isoform a (RBBP4_a, NP_005601.1) but is absent in isoform c (Fig. S4). As SIN3B_3 was observed exclusively within the cytoplasm, this interaction may represent an interaction between cytosolic proteins.

**Figure 4.**
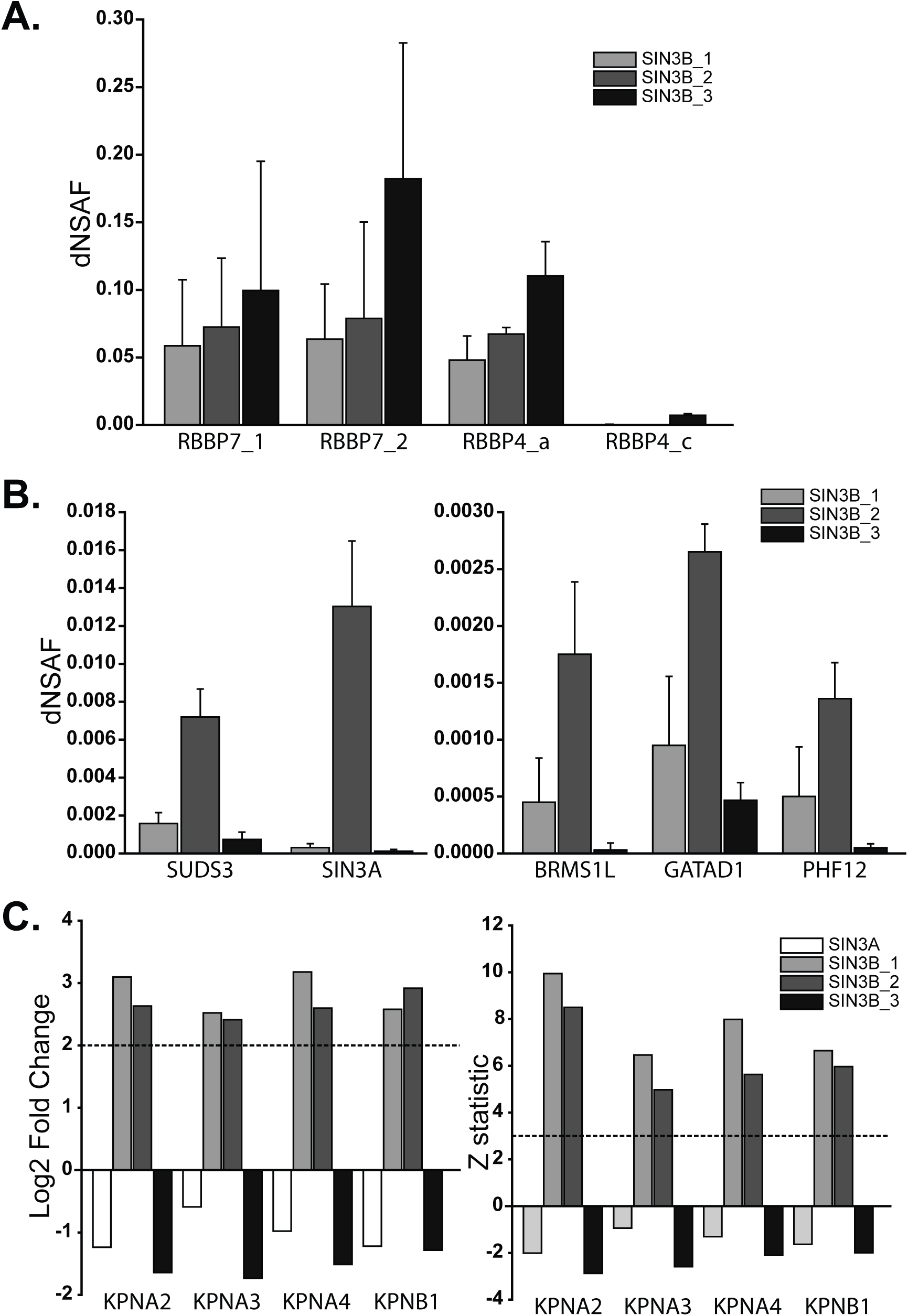
Analysis of interaction partners of SIN3B isoforms. (A) dNSAF values of RBBP4 and RBBP7 proteins identified as being enriched by at least one SIN3B isoform. (B) dNSAF values of SUDS3 and SIN3A (left panel) as well as BRMS1L, GATAD1, and PHF12 (right panel) following SIN3B_1-(light grey), SIN3B_2- (dark grey), and SIN3B_3-(black) purifications (Table S2E). (A-B) *Mean ± S.D., n=*4. (C) log2 fold change (left panel) and Z-statistic (right panel) values calculated by QSPEC v1.3.5 for KPNA2, KPNA3, KPNA4, and KPNB1 in SIN3A-(white), SIN3B isoform 1-(light grey), SIN3B isoform 2-(dark grey), and SIN3B isoform 3-(black) purified samples (**Table S2D**).

Proteins that have homology to Rpd3S- and Rpd3L-specific components also displayed differential enrichment by the SIN3B isoforms (Fig. 4B). Rpd3L-specific component homologs SUDS3 and BRMS1L were both consistently less abundant following SIN3B_1 and SIN3B_3 purifications compared to SIN3B_2 purified samples (Fig. 4B and Table S2E-F). Further, SIN3A met criteria for enrichment by SIN3B_2 (Fig. 1E, Table S2E) but not by SIN3B_1 or SIN3B_3 (Table S2E), suggesting that Sin3 hetero-dimerization is also influenced by the SIN3B HID.

In addition to Rpd3L-specific component homologs, the Rpd3S-specific component homolog PHF12 and its interaction partner GATAD1 were also less abundant in SIN3B_1-purified samples and barely detected in the SIN3B_3 pull-downs compared to SIN3B_2-purified samples (Fig. 4B. Table S2E-F). These data suggests that the HID region likely influences the organization of both major Sin3 interaction networks but is not required for interactions between SIN3B and RBBP4/7.

### SIN3A and SIN3B have divergent nuclear localization signals

While SIN3A, SIN3B_1, and SIN3B_2 were observed within the nucleus, SIN3B_3 was absent from the nucleus. This localization pattern suggests that SIN3B_3 may lack domain(s) required for the nuclear import of this protein. Additionally, SIN3B_1 and SIN3B_2 enriched karyophorins KPNA2, KPNA3, KPNA4, and KPNB1, which are involved in the import of nuclear proteins (Fig. 1E-F, 4C, Table S2E). Notably, SIN3B_1 and SIN3B_2 contain a sequence predicted by cNLS Mapper (25) to be a bipartite nuclear localization signal (NLS) (Fig. 5A, S2). This sequence is absent within SIN3B_3 (Fig. 5A, S2) and this isoform failed to enrich karyophorin proteins (Fig. 4A-B, Table S2E).

**Figure 5.**
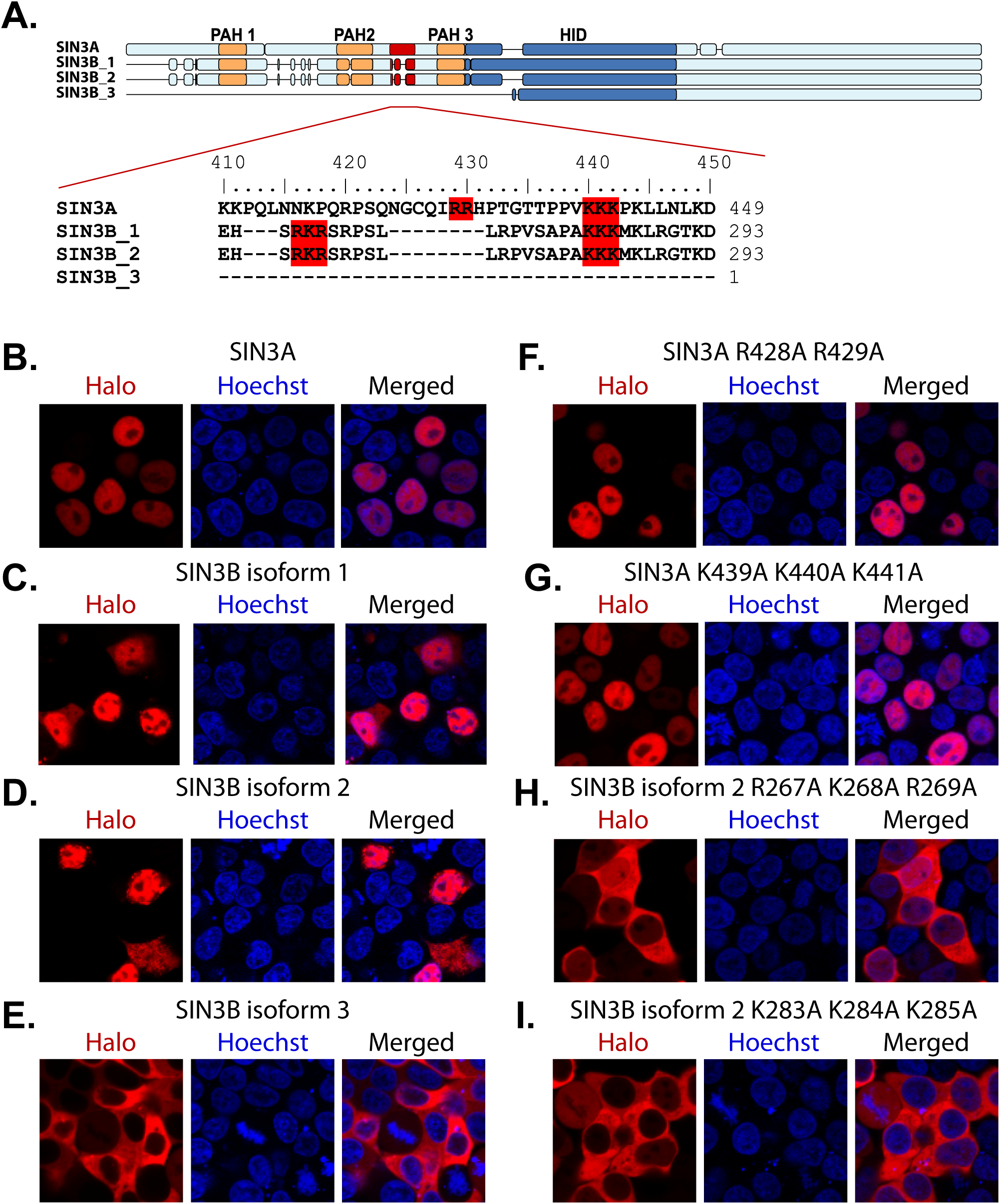
Identification of the SIN3B nuclear localization signal. (A) Alignment of the predicted bipartite nuclear localization sequences in SIN3A and SIN3B. Motifs predicted by cNLS Mapper (25) and residues mutated to test predicted NLS accuracy are highlighted in red. (B-I) Subcellular localization of transiently expressed wild-type (B-E) and mutant (F-I) forms of Halo-tagged recombinant Sin3 proteins in 293T cells. (F-I) Protein names are appended with the identities of mutated residues. HaloTag® TMRDirect™ Ligand and Hoechst 33258 solution to visualize recombinant protein localization (red) and nuclei (blue), respectively.

To test the accuracy of the predicted NLS sequence in SIN3B isoforms, basic residues within the predicted bipartite sequence in SIN3B_2 were mutated to alanine residues. Basic residues found within SIN3A that align with the predicted SIN3B NLS were also mutated. Open reading frames encoding wild-type (Fig. 5B-E) and mutant (Fig. 5F-I) forms of SIN3A and SIN3B proteoforms were transiently expressed in 293T cells as fusions with HaloTag. Mutating residues within either segment of the predicted SIN3B_2 NLS inhibited the nuclear localization the recombinant protein (Fig. 5H-I), indicating that this site functions a bipartite NLS. Surprisingly, the introduction of mutations to homologous residues in SIN3A did not inhibit the nuclear localization of SIN3A (Fig. 5F-G). The observation that KPNA2/3/4 and KPNB1 were not enriched by SIN3A purification (Fig. 1E, 4C, Table S2E) supports the conclusion we derived from the mutational analyses of nuclear localization signals: SIN3A and SIN3B_1/2 are imported to the nucleus via distinct molecular interactions (Figure 6).

**Figure 6.**
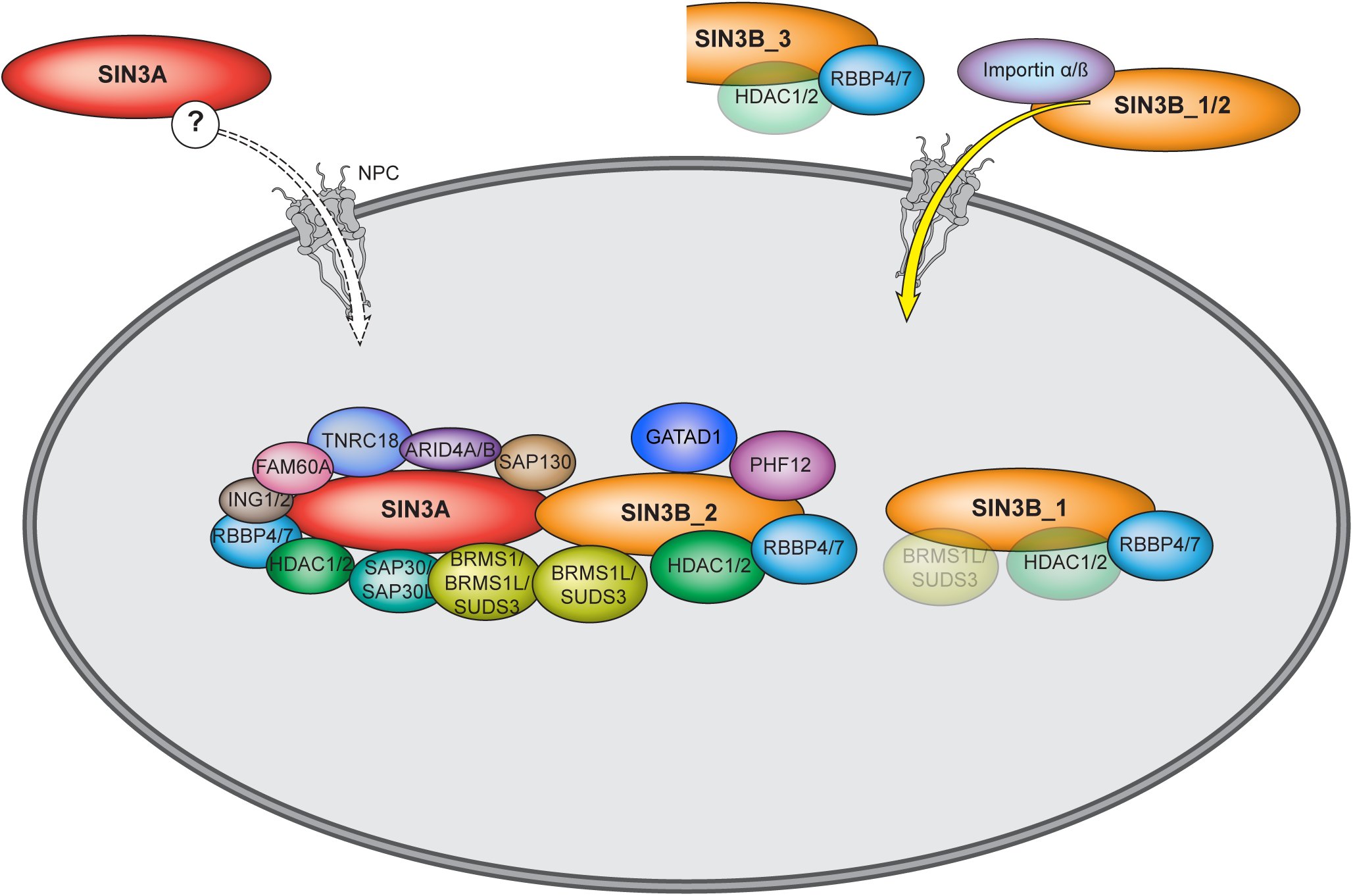
Model of the human SIN3A and SIN3B interactions defined in this study. SIN3A and SIN3B have divergent nuclear localization signals that mediate their nuclear import. SIN3B_3 is not found within the nucleus and has weak interactions with HDAC1/2 and RBBP4/7. SIN3A and SIN3B_2 are capable of forming hetero-oligomeric complex.

## Discussion

To understand the therapeutic potential of targeting Sin3 complex function, we must first characterize the heterogeneous population of Sin3 HDAC complexes. Through a comparative analysis of human Sin3 protein forms, we highlight the influence of paralog switching on complex composition and identify, as well as the shared and unique features of Sin3 protein paralogs.

### SIN3B is a component of two protein interaction networks

The single *S. cerevisiae* Sin3 protein is partitioned into 2 distinct protein complexes, known as Sin3S (or Rpd3S) and Sin3L (or Rpd3L) (5, 6). Common to both yeast complexes is a core of proteins consisting of Sin3, Rpd3, and Ume1, which share homology with human SIN3A/B, HDAC1/2, and RBBP4/7, respectively. While the two yeast Sin3 complexes share a common catalytic subunit, the complexes have distinct subunits that fine-tune complex function and target the complexes to different regions of genes (5, 6). There is mounting evidence that two distinct forms of the Sin3 complex exist in humans and that these complexes can be defined based on subunit homology to yeast Sin3 complex components (26–28). Human SUDS3/BRMS1/BRMS1L, ING1/2, and SAP30/SAP30L have homology with yeast Rpd3L-subunits Sds3, Pho23, and Sap30, respectively. Additionally, human PHF12 and MORF4L1 share homology with yeast Rpd3S-specific subunits Rco1 and Eaf3.

Unlike yeast, humans have two gene paralogs, *SIN3A* and *SIN3B*, that encode Sin3 proteins. To determine if the identity of the Sin3 paralog within a complex defines which interaction network is formed, we performed a comparative analysis of SIN3A and SIN3B protein-protein interactions. Our results suggest that both SIN3A and SIN3B can interact with HDAC1/2 and RBBP4/7, proteins with homology to yeast core proteins found within both Rpd3L and Rpd3S complexes. SIN3B was found to interact with SUDS3/BRMS1L and PHF12. These results show that, much like yeast Sin3, human SIN3B can interact with homologs of both Rpd3S and Rpd3L complex-specific components. Though PHF12 was initially identified as an interaction partner of SIN3A (29, 30), SIN3A-purified samples were devoid of peptides mapping to PHF12. Thus, SIN3A enriched for homologs of Rpd3L-specific but not Rpd3S-specific components. Thus, Sin3 protein identity influences Sin3 complex composition and confirm previous reports that PHF12 interacts with SIN3B but not SIN3A (31).

An interesting finding was that SIN3B_2 has the capacity to enrich SIN3A, which suggest hetero-oligomeric forms of human Sin3 complexes exist in humans. SUDS3, BRMS1, and BRMS1L have been proposed to induce dimerization of Sin3 complexes (32). It remains to be determined if the formation of SIN3A-SIN3B hetero-oligomers requires the presence of SUDS3/BRMS1/BRMS1L or if SIN3A and SIN3B directly interact with one another.

In addition to proteins with clear homology to yeast Sin3 complex components, several human Sin3 interaction partners exist that have no clear homology to yeast proteins. One such protein GATAD1, a known Sin3 interaction network component (26, 27), was enriched by SIN3B_2 but not SIN3A (Fig. 1E-F). TNRC18, FAM60A, SAP130, and ARID4A/B are additional Sin3 interaction partners (33, 34) that met criteria for enrichment by SIN3A but not SIN3B_2. As ARID4A/B have proposed DNA binding capacities (35), such divergent behavior by SIN3A and SIN3B likely has functional consequences. It is important to note that peptides mapping to these proteins were observed following SIN3B_2 purification but that these proteins did not meet criteria for enrichment (Fig. 1F, Table S2E). Thus, interactions between SIN3B and these proteins may be weak or indirect. As SIN3A was enriched by SIN3B_2, a possible explanation for the weak identification of these protein following SIN3B_2 purification is that indirect interactions between SIN3B_2 and FAM60A, TNRC18, ARID4A/B, and SAP130 could be mediated by SIN3A (Fig. 6).

### Protein domain organization is partially conserved between SIN3A and SIN3B

SIN3A contains several experimentally defined domains. A region of mouse SIN3A that is sufficient and necessary for interactions with HDAC1/2 was previously identified and termed the HDAC interaction domain, or HID (24, 32, 36). Additionally, 3 PAH domains are present in the N-terminal half of SIN3A and there is evidence which suggests these domains mediate interaction between SIN3A and a multitude of transcription factors (4, 37). Few studies of SIN3B domain organization have been performed. However, several regions have high homology to defined domains within SIN3A. While a HID has not been experimentally defined within SIN3B, an alignment of the mouse SIN3A and SIN3B protein sequences shows that residues 388 to 651 of SIN3B_2 have approximately 88% sequence similarity to residues 545 to 808 in SIN3A.

While isoform 2 (NP_001284524) likely represents the principal SIN3B isoform in humans, alternative splicing produces multiple SIN3B isoforms in mouse (38, 39) and it was reported that ratios of human *SIN3B* gene products are responsive to cellular queues (38). Though alternative isoforms of SIN3B are likely a small portion of the total SIN3B within humans, these alternative proteoforms provide insight into the domain organization of SIN3B. SIN3B_1 has an additional exon within a region that has high homology to the SIN3A HID (Fig 2A, S2). Therefore, we used this alternative SIN3B proteoform to provide insight into the dependence of SIN3B protein-protein interaction on the SIN3B HID. We demonstrate that isoform 1, which contains a 32-residue sequence not found within isoform 2, has a decreased catalytic potential as well as weaker associations with Sin3 network components (Fig. 2, Table S2). It should be noted that isoform 1 was still capable of enriching HDAC1/2 (Table S2). Thus, the addition of this exon does not completely abolish interactions with HDAC1/2 but does diminish both HDAC1/2 binding and the complex’s catalytic capacity (Fig. 3C, 3E).

A region within SIN3B that has sequence homology to a portion of the SIN3A HID is critical for interactions between SIN3B and PHF12 (31). Our results support this observation as the additional exon found within SIN3B_1 disrupts interactions between SIN3B and PHF12 (Fig. 5F). Additionally, we observed poor interactions between SIN3B_1 and homologs of yeast Sin3L components SUDS3 and BRMS1L (Fig. 3A, 4B, Table S2). Results obtained by others suggest that the HID regions is critical for interactions between SIN3A and SUDS3 (21). Together, these results suggest that the organizing role of the HID region is conserved between SIN3A and SIN3B. As SUDS3/BRMS1 and PHF12 were less abundant in SIN3B_1 compared to SIN3B_2 enrichments, the HID is likely important for the organization of both forms of the Sin3 complex in humans.

Notably, SIN3A was less abundant following SIN3B_1 enrichment compared to SIN3B_2 enrichment (Fig. 4B, Table S2E). Thus, the HID appears to be critical for hetero-oligomerization of the Sin3 complex. Previous findings suggest that yeast Sds3 is essential for complex integrity (17) and mammalian BRMS1 (32) and SUDS3 (21, 32) are capable of forming dimers. As SUDS3 and BRMS1L were enriched by both SIN3A and SIN3B_2, it is possible that these proteins mediate the formation of SIN3A-SIN3B_2 hetero-oligomeric complexes.

SIN3B_3 provides us with insight into the mechanisms responsible for SIN3B nuclear import. This isoform, resulting from an alternative start codon, is significantly shorter than SIN3B_1/SIN3B_2 and has absent or disrupted PAH and HID domains (Fig. 2A). Interestingly, recombinant SIN3B_3 failed to localize to the nucleus (Fig. 2C). This isoform lacks a predicted NLS signal and prompted us to investigate the requirement for this sequence for SIN3B nuclear import. Analysis of the SIN3B_3 interaction network revealed that, unlike other SIN3B isoforms, SIN3B_3 did not enrich KPNA2/3/4 or KPNB1. The introduction of mutations to the predicted NLS of SIN3B isoform 2 resulted in cytoplasmic localization; however, mutations to conserved or similar residues within SIN3A had no effect on nuclear localization. Thus, our findings indicate that SIN3A and SIN3B have distinct nuclear localization signals. Consistently, it has recently been shown that nonsense mutations at residue 949 in SIN3A results in a truncated protein with cytoplasmic localization (40), suggesting that the NLS of SIN3A resides within its C-terminus and is distinct from SIN3B NLS. Additionally, SIN3A, unlike SIN3B_2, did not enrich KPNA2 KPNA3, KPNA4, or KPNB1 (Fig. 4C, Table S2E), further suggesting that SIN3A and SIN3B may experience unique modes of nuclear import (Fig. 6).

Not all identified proteins displayed differential interactions with SIN3B isoforms. In fact, RBBP4 and RBBP7 were consistently identified as the most abundant non-bait proteins in all SIN3B isoform purifications (Fig. 4A). As SIN3B_1 and SIN3B_3 enriched RBBP4/7, these results show that the C-terminal half of SIN3B is sufficient for interactions with these proteins. While the HID may act as an organizing domain for much of the complex, RBBP4/7 do not depend on this region for interactions with SIN3B. Thus, while HDAC1/2 and RBBP4/7 likely form a shared core complex with Sin3 proteins in humans, they likely do so via interaction with distinct regions of Sin3 proteins. This is surprising as HDAC1/2 and RBBP4/7 co-exist within multiple HDAC complexes and have been proposed to form a preformed sub-module (41).

In total, these results provide insight into the shared and unique properties of human Sin3 scaffolding proteins. While a conserved HID is present in both proteins, the identity of the Sin3 protein found within complexes influences their interaction networks and, likely, composition. We also show that hetero-oligomeric forms of the Sin3 complex can exist as SIN3B enriches SIN3A. These findings highlight the influence of paralog switching on protein complex composition and outline the need for future studies that further delineate the unique functions of the distinct classes of Sin3 complexes. Future studies should take into account additional heterogeneity within population of Sin3 complexes that is introduced by other complex subunits that exist as protein paralogs.

## Abbreviations

AP: affinity purification
APMS: affinity purification mass spectrometry
DMSO: dimethyl sulfoxide
dNSAF: distributed normalized spectral abundance factor
dS: distributed spectral counts
FBS: fetal bovine serum
FDA: Food and Drug Administration
FDR: false discovery rate
HDAC: histone deacetylase
HDACi: HDAC inhibitor
HID: HDAC interaction domain
HPLC: high performance liquid chromatography
KDRI: Kazusa DNA Research Institute
MudPIT: multidimensional protein identification technology
NLS: nuclear localization signal
PAH: paired amphipathic helix
RBBP4_a: RBBP4 isoform a
RBBP4_c: RBBP4 isoform c
RFU: relative fluorescence units
Rpd3L (Sin3L): Sin3 Large
Rpd3S (Sin3S): Sin3 Small
SAHA: suberoylanilide hydroxamic acid
SIN3B_1: SIN3B isoform 1
SIN3B_2: SIN3B isoform 2
SIN3B_3: SIN3B isoform 3

## Acknowledgments

Original data underlying this manuscript can be accessed from the Stowers Original Data Repository at http://www.stowers.org/research/publications/libpb-1445. Research reported in this publication was supported by the Stowers Institute for Medical Research and the National Institute of General Medical Sciences of the National Institutes of Health under Award Number F32GM122215 (M.K.A.), F31GM131536 (C.G.K), and R01GM112639 (M.P.W.). The content is solely the responsibility of the authors and does not necessarily represent the official views of the National Institutes of Health.

## Author contributions

M.K.A., C.A.S.B., J.L.T., and M.P.W. designed research; M.K.A., C.A.S.B., J.L.T., M.K., and C.G.K. performed research; M.K.A., C.A.S.B., J.L.T., C.G.K., and M.P.W. contributed new reagents/analytic tools; M.K.A., C.A.S.B., J.L.T., M.E.S., M.K., C.G.K., L.F. and M.P.W. analyzed data; M.K.A. and M.P.W. wrote the paper.

## Experimental Procedures

### Preparation of expression vectors and expression in Flp-In™-293 cell lines

To prepare expression constructs for SIN3A expression, the V109A point mutation present in Kazusa DNA Research Institute (KDRI) clone # FHC11647 was corrected using targeted mutagenesis. SIN3A was then excised with AsiSI and PmeI then into cloned into AsiSI and Eco53KI sites of pFC14A (Promega Corporation, Madison, WI) to add a C-terminal HaloTag®. SIN3A with an in-frame C-terminal HaloTag® was excised and cloned into MluI and NotI sites of pcDNA™5/FRT Mammalian Expression Vector (Thermo Fisher Scientific, Waltham, MA). To prepare expression constructs for SIN3B isoform 2 (SIN3B_2) expression, KDRI clone # FHC01991 was digested with AsiSI and PmeI, then cloned into AsiSI and Eco53KI sites of pFC14A to add a C-terminal HaloTag®. SIN3B_2 with an in-frame C-terminal HaloTag® was excised and cloned into MluI and NotI sites of pcDNA™5/FRT Mammalian Expression Vector. To prepare SIN3B isoform 1 (SIN3B_1) expression constructs, the MiniGene™ (Integrated DNA Technologies, Coralville, Iowa) described in Fig. S5A was digested with NsiI and BsiWI and cloned into these restriction sites of KDRI clone # FHC01991. This construct was then digested with AsiSI and BsiWI then cloned into AsiSI and BsiWI sites of the above described SIN3B_2 in pFC14A to create SIN3B_1 with a C-terminal HaloTag®. SIN3B_1 with an in-frame C-terminal HaloTag® was excised and cloned into MluI and NotI sites sites of pcDNA™5/FRT Mammalian Expression Vector. To prepare SIN3B isoform 3 (SIN3B_3) expression constructs, a gBlock® (Integrated DNA Technologies), described in Fig. S5B was digested with AsiSI and BsiWI then cloned into the above described SIN3B_2-pFC14A. SIN3B_3 with an in-frame C-terminal HaloTag® was excised and cloned into MluI and NotI sites of pcDNA™5/FRT Mammalian Expression Vector.

Stable cell lines were produced using Flp-In™-293 cells (Thermo Fisher Scientific), authenticated by STR profiling (FTA barcode: STR14169), and tested for mycoplasma using mycoplasma detection kits (American Type Culture Collection, Manassas, VA). The day before transfection, cells were plated at 50% confluency (DMEM/10% FBS) onto a 100 mm tissue culture plate and incubated at 37 °C, 5% CO2 overnight. The following day, the plate was washed two times with Opti-MEM, then incubated until ready to use with 8 mL Opti-MEM with GlutaMAX supplement (Thermo Fisher Scientific). Plasmid DNA (4 µg total; 3.6 µg pOG44 + 0.4 µg DNA of interest) was added to 800 µl of Opti-MEM with GlutaMAX supplement along with 16 µl FuGENE® HD Transfection Reagent (Promega Corporation), incubated for 15-30 minutes in a biosafety cabinet, then added dropwise to the prepared plate. One mL of FBS (Peak Serum, Inc, Wellington CO) was added the next morning. On day three of incubation, cells were split 1:10 and placed into selection media (DMEM/10% FBS/100 µg/mL Hygromycin B). Media was changed every three days for a total of three media changes. After two weeks, colonies were visible and picked for screening.

### Fluorescence Microscopy

Flp-In™-293 cell lines stably expressing HaloTag® fusion proteins were seeded at 40% confluency in 35 mm MatTek glass bottom dishes (MatTek Corporation, Ashland, MA) containing DMEM supplemented with penicillin-streptomycin solution, GlutaMAX supplement, and FBS to a final concentration of 10%. Cell media was supplemented with HaloTag® TMRDirect™ Ligand (Promega Corporation) 16-24 h after seeding to a final concentration of 20 nM and cells were incubated for an additional 16-24 h. Hoechst 33258 solution (Sigma Aldrich Corporation, St. Louis, MO) was added 80 min prior to imaging cells. Cells were washed two times with Opti-MEM media and imaged in Opti-MEM media. Images were captured on a PerkinElmer Life Sciences UltraVIEW VoX spinning disk microscope (PerkinElmer, Inc. Waltham, MA), Axiovert 200M base (Carl Zeiss AG, Oberkochen, Germany), or an inverted LSM-700 point scanning confocal microscope controlled by Zeiss Zen software (Carl Zeiss AG). A 40X plan-apochromat (NA 1.4) oil objective was used to acquire images when operating the LSM-700 microscope. Detection wavelength ranges were 300-483 nm for Hoechst and 570-800 nm for HaloTag® TMRDirect™ Ligand. A SP 490 filter set and LP 490 filter set were employed when imaging Hoechst and HaloTag® TMRDirect™ Ligand, respectively on the LSM-700 microscope.

Media conditions for transient transfection of 293T cells (American Type Culture Collection) with plasmid DNA were as stated for the imaging of the stable cell lines. Cells continued to grow 16-24 h after seeding at 40% in 35 mm MatTek glass bottom dishes before transfection. Cells were transfected with Opti-MEM media containing 2.5 µg of plasmid, 5 µL Lipofectamine LTX along with 2.5 µg PLUS Reagent (Thermo Fisher Scientific), and 20 nM final concentration HaloTag® TMRDirect™ Ligand. After 16-24 h of incubation at 37 °C and 5% CO_2_, Hoechst 33258 solution was added to the dishes and incubation was continued for 1 h. Cells were washed two times with Opti-MEM media before imaging in Opti-MEM media.

### Affinity purification of recombinant proteins from Flp-In™-293 cells and Multidimensional Protein Identification Technology (MudPIT) Analysis of Transiently Expressed Protein

Cells were lysed and recombinant proteins were isolated using Magne® HaloTag® Beads (Promega) as previously described (15). Affinity purified (AP) proteins were TCA precipitated, digested with Lys-C or rLys-C, Mass Spec Grade (Promega Corporation) then Sequencing Grade Trypsin (Promega Corporation). Peptides were loaded onto triphasic MudPIT microcapillary columns as previously (42). Columns were placed in-line with an 1100 Series HPLC system (Agilent Technologies, Inc., Santa Clara, CA) coupled to a linear ion trap mass spectrometer (Thermo Fisher Scientific) and peptides were resolved using 10-step MudPIT chromatography as previously described (43).

Acquired .RAW files were converted to .ms2 files using RAWDistiller (44). ProLuCID v1.3.5 (45) was used to match spectra against a database containing human protein sequences (National Center of Biotechnology Information, June 2016 release) along with shuffled sequences for false discovery rate (FDR) estimation. The database was searched for fully tryptic peptides with static modification of +57 daltons for cysteine and a dynamic modification of +16 daltons for methionine residues. DTASelect and Contrast (46) were used to filter results and NSAFv7 (47) was used to calculate label-free quantitative dNSAF values and generate final reports (Tables S2A-B). The spectral FDR mean ± s.d. for the 19 MudPIT runs was 0.198% ± 0.105%, the mean peptide FDR was 0.355% ± 0.225%, and the mean ± s.d protein FDR was 1.30% ± 0.769%. A DTASelect filter also established a minimum peptide length of 7 amino acids, and proteins that were subsets of others were removed using the parsimony option in Contrast.

Data that has been previously described was included in our analyses and is summarized in Table S1. All mass spectrometry data has been deposited into the MassIVE repository (http://massive.ucsd.edu). Data set identifiers are supplied in Table S1.

### Statistical analysis of proteomics data sets

A minimum of three biological replicates were acquired for each affinity purification mass spectrometry (APMS) analysis. To identify high-confidence interaction partners, QSPEC v1.3.5 (16) was used to calculate Z-statistic, log2 fold change, and FDR values of identified proteins. Prey proteins that were not present in at least half of at least one bait protein purification (Table S2C) were excluded prior to QSPEC scoring (Tables S2D). QSPEC analysis was performed with a burn in value of 2000 and 10,000 iterations. To identify enriched proteins over negative AP controls, Z-statistic values of ≥ 3, log2 fold change values ≥ 2 were selected as filter values. Only proteins with FDR values ≤ 0.05 were considered for analysis. Clustered heatmaps of normalized dNSAF values were generated using the python package Seaborn and the unweighted pair group method with arithmetic mean. Normalized dNSAF values were obtained by subtracting the minimum dNSAF value and dividing by the maximum dNSAF value. The Circos plot (48) of SIN3A and SIN3B_2 interactions was generated using average dNSAF values for enriched by SIN3A and/or SIN3B_2.

### Enzyme activity assays

HDAC activity assays of transiently produced proteins were performed as described (49). Briefly, approximately 1 × 10^7^ 293T cells were plated in 150 mm dishes and cultured in 25 mL DMEM + 10% fetal bovine serum + 1x GlutaMAX Supplement. 24 h after seeding, cells were transfected with 7.5 µg plasmid DNA, 7.5 µL Plus Reagent, and 50 µL Lipofectamine LTX diluted in 6.6 mL OptiMEM. Cells were harvested after an additional 48 h of culture. Two mg of whole cell extract was added to 100 µL of washed Magne® HaloTag® Beads slurry and incubated at 4 °C for 2 h. Beads were washed 4 times with 1 mL cold TBS pH 7.4 + 0.05% Igepal CA-630 (Sigma Aldrich Corporation). Protein was eluted with 5 units AcTEV™ Protease (Thermo Fisher Scientific) in 100 µL of 50 mM Tris-HCl pH 8.0, 0.5 mM EDTA, 1 mM DTT for 16 h at 4 °C. The HDAC activity assay was based on previously published procedures (50, 51). In brief, 10 µL of the 100 µL of the purified protein was diluted with 32.5 µL TBS (25 mM Tris, 150 mM NaCl, 2mM KCl, pH 7.4). Samples were supplemented with 2.5 µL of DMSO or 200 µM SAHA (Cayman Chemical Company, Ann Arbor, MI) resuspended in DMSO for a final concentration of 10 µM SAHA. 5 µL of 1 mM Boc-Lys(Ac)-AMC (APExBIO Technology LLC, Houston, TX) was added to each reaction to a final concentration of 100 µM. Final reaction volume was 50 µL. Reactions were incubated at 37 °C for 1 h. Reactions were quenched with 2.5 µl of 200 µM SAHA and incubated at 37 °C for 5 min. 6 µL of 50 mg/mL trypsin was added to reaction for a final concentration of 5 mg/mL. Reactions were incubated an additional 1 h at 37 °C. Fluorescence was measured with a SPECTRAmax GEMINI XS (Molecular Devices, San Jose, CA) using an excitation wavelength of 355, emission wavelength of 460 nm, and a cutoff wavelength of 455 nm.

### Sequence Alignment

Graphic alignment of SIN3A (NP_001138829.1), SIN3B_1 (NP_056075.1), SIN3B_2 (NP_001284524.1), and SIN3B_3 (NP_001284526.1) was generated using ETE v3 (52) and ClustalO (53). Pairwise alignments were generated using the EMBOSS-Needle algorithm (54).

## Figures

**Supplementary Figure S1 Related to Figure 1.** Pairwise sequence alignment of SIN3A, (NP_001138829.1) and SIN3B isoform 2 (NP_001284524.1) generated using EMBOSS-Needle (54).

**Supplementary Figure S2 Related to Figure 2.** Protein alignment of SIN3A and SIN3B isoforms generated with ClustalO (53). A sequence within SIN3B_1 and SIN3B_2 that possesses a cNLS Mapper score of 12.5 as a bipartite NLS is highlighted in yellow. A sequence within SIN3A possesses a cNLS Mapper score of 4.0 as a bipartite NLS is also highlighted in yellow. Residues within human SIN3A that are homologous to the mouse SIN3A HID (24) are in red. **Supplementary Figure S3 Related to Figure 3D-E.** Image of Western blot shown in **Fig. 3D** with intensity values displayed.

**Supplementary Figure S4 Related to Figure 4A.** Pairwise sequence alignment of RBBP4 isoform a, (NP_005601.1) and RBBP4 isoform c (NP_001128728.1) generated using EMBOSS-Needle (54). A sequence within RBBP4 isoform a that possesses a cNLS Mapper score of 3.8 as a bipartite NLS is highlighted in yellow.

**Supplementary Figure S5 Related to Figures 2, 3, 4, 5.** (A) The MiniGene™ (Integrated DNA Technologies) sequence used to create SIN3B isoform 1. NsiI and BsiWI recognition sequences are underlined. (B) The gBlock® (Integrated DNA Technologies) sequence used to create SIN3B isoform 3. AsiSI and BsiWI recognition sequences are underlined.

**Supplementary Table S1 Related to Figures 1, 2, 3, 4, 5.** Data Availability

**Supplementary Table S2 Related to Figures 1, 2, 3, and 4**. Identification, label-free quantitation, and statistical analysis of proteins detected in APMS analyses of SIN3 paralogs and isoforms

**Supplementary Table S3 Related to Figures 3.** HDAC activity assay.

