## Supplemental Figures for "Differential complex formation via paralogs in the human Sin3 protein interaction network"

### Supplementary Figure S1 | Adams et al.

**Needle Pairwise Sequence Alignment**

```
Protein          Accession
SIN3A:           NP_001138829.1
SIN3B isoform 2: NP_001284524.1

Program: needle
        -datafile EBLOSUM62
        -gapopen 10.0
        -gapextend 0.5
        -endopen 10.0
        -endextend 0.5
        -Matrix: EBLOSUM62
        -Gap_penalty: 10.0
        -Extend_penalty: 0.5

Length: 1295
Identity:     626/1295 (48.3%)
Similarity:   812/1295 (62.7%)
Gaps:         187/1295 (14.4%)
Score: 3068.0

NP_001138829.      1 MKRRLDDQESPVYAAQQRRIPGSTEAFPHQHRVLAPAPPVYEAVSETMQS     50

NP_001284524.      1 --------------------------------------------------      0

NP_001138829.     51 ATGIQYSVTPSYQVSAMPQS------SGSHGPAIAAVHSSHHHPTAVQPH     94
                                     |..:      ||:.|||...:..:....:.    
NP_001284524.      1 ----------------MAHAGGGSGGSGAGGPAGRGLSGARWGRSG----     30

NP_001138829.     95 GGQVVQSHAHPAPPVAPVQGQQQFQRLKVEDALSYLDQVKLQFGSQPQVY    144
                           |..|...||            .|||||:||||||::|||.|..|
NP_001284524.     31 ------SAGHEKLPV------------HVEDALTYLDQVKIRFGSDPATY     62

NP_001138829.    145 NDFLDIMKEFKSQSIDTPGVISRVSQLFKGHPDLIMGFNTFLPPGYKIEV    194
                     |.||:|||||||||||||||.||||||..|||||:|||.|||.||:|::
NP_001284524.     63 NGFLEIMKEFKSQSIDTPGVIRRVSQLFHEHPDLIVGFNAFLPLGYRIDI    112

NP_001138829.    195 QTNDMVNVTTPGQVHQIPTHGIQPQPQPPPQHPSQPSAQSAPAPAQPAPQ    244
                     ..|..:|:.:|                    ..||.::.:....|:...|
NP_001284524.    113 PKNGKLNIQSP--------------------LTSQENSHNHGDGAEDFKQ    142

NP_001138829.    245 PPPAKVSKPSQLQAHTPASQQTPPLPPYASPRSPPVQPHTPVTISLGTAP    294
                     ..|.|..||           |.|                           
NP_001284524.    143 QVPYKEDKP-----------QVP---------------------------    154

NP_001138829.    295 SLQNNQPVEFNHAINYVNKIKNRFQGQPDIYKAFLEILHTYQKEQRNAKE    344
                      |:::. ||||:||:||||||.||...|:||::||||||||||||.|.: 
NP_001284524.    155 -LESDS-VEFNNAISYVNKIKTRFLDHPEIYRSFLEILHTYQKEQLNTR-    201

NP_001138829.    345 AGGNYTPALTEQEVYAQVARLFKNQEDLLSEFGQFLPDANSSVLLSKTTA    394
                       |.....::|:||:.:||.||:.|||||||||||||:|..|:.......
NP_001284524.    202 --GRPFRGMSEEEVFTEVANLFRGQEDLLSEFGQFLPEAKRSLFTGNGPC    249

NP_001138829.    395 EKVDSVRNDHGGTVKKPQLNNKPQRPSQNGCQIRRHPTGTTPPVKKKPKL    444
                     |.....:|:|.   |.|:.:.|..|||    .:|    ..:.|.|||.||
NP_001284524.    250 EMHSVQKNEHD---KTPEHSRKRSRPS----LLR----PVSAPAKKKMKL    288

NP_001138829.    445 LNLKDSSMADASKHGGGTESLFFDKVRKALRSAEAYENFLRCLVIFNQEV    494
                     ...||.|:|...|:|...|..||||||:.|:|.|.|||||||:.:||||:
NP_001284524.    289 RGTKDLSIAAVGKYGTLQEFSFFDKVRRVLKSQEVYENFLRCIALFNQEL    338
```

### Supplementary Figure S1 | Adams et al.

```
NP_001138829.    495 ISRAELVQLVSPFLGKFPELFNWFKNFLGYKESVHLETYP--KERATEGI    542
                     :|.:||:||||||||||||||..||:|||.||   |...|  .:|:.:||
NP_001284524.    339 VSGSELLQLVSPFLGKFPELFAQFKSFLGVKE---LSFAPPMSDRSGDGI    385

NP_001138829.    543 AMEIDYASCKRLGSSYRALPKSYQQPKCTGRTPLCKEVLNDTWVSFPSWS    592
                     :.|||||||||:|||||||||:||||||:|||.:||||||||||||||||
NP_001284524.    386 SREIDYASCKRIGSSYRALPKTYQQPKCSGRTAICKEVLNDTWVSFPSWS    435

NP_001138829.    593 EDSTFVSSKKTQYEEHIYRCEDERFELDVVLETNLATIRVLEAIQKKLSR    642
                     |||||||||||.|||.::||||||||||||||||||||||||::||||||
NP_001284524.    436 EDSTFVSSKKTPYEEQLHRCEDERFELDVVLETNLATIRVLESVQKKLSR    485

NP_001138829.    643 LSAEEQAKFRLDNTLGGTSEVIHRKALQRIYADKAADIIDGLRKNPSIAV    692
                     ::.|:|.|||||::||||||||.|:|:.|||.|||.:||:.|:|||..||
NP_001284524.    486 MAPEDQEKFRLDDSLGGTSEVIQRRAIYRIYGDKAPEIIESLKKNPVTAV    535

NP_001138829.    693 PIVLKRLKMKEEEWREAQRGFNKVWREQNEKYYLKSLDHQGINFKQNDTK    742
                     |:||||||.|||||||||:||||:||||.||.||||||||.:||||||||
NP_001284524.    536 PVVLKRLKAKEEEWREAQQGFNKIWREQYEKAYLKSLDHQAVNFKQNDTK    585

NP_001138829.    743 VLRSKSLLNEIESIYDERQEQATEENAGVPVGPHLSLAYEDKQILEDAAA    792
                     .||||||||||||:|||.|||.:|..:.....|||...|||:||||||||
NP_001284524.    586 ALRSKSLLNEIESVYDEHQEQHSEGRSAPSSEPHLIFVYEDRQILEDAAA    635

NP_001138829.    793 LIIHHVKRQTGIQKEDKYKIKQIMHHFIPDLLFAQRGDLSDVEEEEEEEM    842
                     ||.::||||..|||||:..|.|::|.|:|.|.|:|:.||...||..:|:.
NP_001284524.    636 LISYYVKRQPAIQKEDQGTIHQLLHQFVPSLFFSQQLDLGASEESADEDR    685

NP_001138829.    843 D-----VDEATGAVKKHNGVGGSPPKSKLLFSNTAAQKL---------RG    878
                     |     ..:.:...|...|...|||:.|..|.:..|.:.         :.
NP_001284524.    686 DSPQGQTTDPSERKKPAPGPHSSPPEEKGAFGDAPATEQPPLPPPAPHKP    735

NP_001138829.    879 MDEVYNLFYVNNNWYIFMRLHQILCLRLLRICSQAERQIEEENREREWER    928
                     :|:||:||:.|||||.|:||||.||.|||:|..||::|:.|...|:|.|:
NP_001284524.    736 LDDVYSLFFANNNWYFFLRLHQTLCSRLLKIYRQAQKQLLEYRTEKEREK    785

NP_001138829.    929 EVLGIKRDKSDSPAIQLRLKEPMDVDVEDYYPAFLDMVRSLLDGNIDSSQ    978
                     .:...:|:|...||::||||:|.:|::|:|||||||||||||:|:||.:|
NP_001284524.    786 LLCEGRREKGSDPAMELRLKQPSEVELEEYYPAFLDMVRSLLEGSIDPTQ    835

NP_001138829.    979 YEDSLREMFTIHAYIAFTMDKLIQSIVRQLQHIVSDEICVQVTDLYLAEN   1028
                     |||:||||||||||:.||||||:|:|.|||.|:|||::|::|.:|||.|.
NP_001284524.    836 YEDTLREMFTIHAYVGFTMDKLVQNIARQLHHLVSDDVCLKVVELYLNEK    885

NP_001138829.   1029 NNGATGGQLNTQNSRSLLESTYQRKAEQLMSDENCFKLMFIQSQGQVQLT   1078
                     ..||.||.|:::..|:..|::||.|||:.|:||||||:||:|.:|||.:|
NP_001284524.    886 KRGAAGGNLSSRCVRAARETSYQWKAERCMADENCFKVMFLQRKGQVIMT    935

NP_001138829.   1079 IELLDTEEENSDDPVEAERWSDYVERYMNSDTTSPELREHLAQKPVFLPR   1128
                     ||||||||..::||||.:..:.|||:|:.::..|....|....|||||.|
NP_001284524.    936 IELLDTEEAQTEDPVEVQHLARYVEQYVGTEGASSSPTEGFLLKPVFLQR    985

NP_001138829.   1129 NLRRIRKCQRGREQQEKEGKEGNSKKTMENVDSLDKLECRFKLNSYKMVY   1178
                     ||::.|:..:..:.:...|:..:|.|.:..|:|...::|||||:::|||:
NP_001284524.    986 NLKKFRRRWQSEQARALRGEARSSWKRLVGVESACDVDCRFKLSTHKMVF   1035

NP_001138829.   1179 VIKSEDYMYRRTALLRAHQSHERVSKRLHQRFQAWVDKWTKEHVPREMAA   1228
                     ::.||||||||..|.||.|....|..|.||.|:.|..:|.:::|..|.|:
NP_001284524.   1036 IVNSEDYMYRRGTLCRAKQVQPLVLLRHHQHFEEWHSRWLEDNVTVEAAS   1085

NP_001138829.   1229 ETSKWLMGEGLEGLVPCTTTCDTETLHFVSINKYRVKYGTVFKAP   1273
                     ....|||||..|.:|||.|.|:|..:|.:.:.:|||:|.....:|
NP_001284524.   1086 LVQDWLMGEEDEDMVPCKTLCETVHVHGLPVTRYRVQYSRRPASP   1130
```

### Supplementary Figure S2 | Adams et al.

```
SIN3A:              NP_001138829.1
SIN3B isoform 1:    NP_056075.1
SIN3B isoform 2:    NP_001284524.1
SIN3B isoform 3:    NP_001284526.1

NP_001138829.1      MKRRLDDQESPVYAAQQRRIPGSTEAFPHQHRVLAPAPPVYEAVSETMQSATGIQYSVTP      60
NP_001284526.1      ------------------------------------------------------------      0
NP_056075.1         ------------------------------------------------------------      0
NP_001284524.1      ------------------------------------------------------------      0


NP_001138829.1      SYQVSAMPQSSGSHGPAIAAVHSSHHHPTAVQPHGGQVVQSHAHPAPPVAPVQGQQQFQR      120
NP_001284526.1      ------------------------------------------------------------      0
NP_056075.1         ------MAHAGGGSGGSG-----------AGGPAGRGLSGARW----G-RSGSAGHEKLP      38
NP_001284524.1      ------MAHAGGGSGGSG-----------AGGPAGRGLSGARW----G-RSGSAGHEKLP      38


NP_001138829.1      LKVEDALSYLDQVKLQFGSQPQVYNDFLDIMKEFKSQSIDTPGVISRVSQLFKGHPDLIM      180
NP_001284526.1      ------------------------------------------------------------      0
NP_056075.1         VHVEDALTYLDQVKIRFGSDPATYNGFLEIMKEFKSQSIDTPGVIRRVSQLFHEHPDLIV      98
NP_001284524.1      VHVEDALTYLDQVKIRFGSDPATYNGFLEIMKEFKSQSIDTPGVIRRVSQLFHEHPDLIV      98


NP_001138829.1      GFNTFLPPGYKIEVQTNDMVNVTTPGQVHQIPT-HGIQPQPQPPPQHPSQPSAQSAPAPA      239
NP_001284526.1      ------------------------------------------------------------      0
NP_056075.1         GFNAFLPLGYRIDIPKNGKLNIQSPLTSQENSHNHGD-----------------GA----      137
NP_001284524.1      GFNAFLPLGYRIDIPKNGKLNIQSPLTSQENSHNHGD-----------------GA----      137


NP_001138829.1      QPAPQPPPAKVSKPSQLQAHTPASQQTPPLPPYASPRSPPVQPHTPVTISLGTAPSLQNN      299
NP_001284526.1      ------------------------------------------------------------      0
NP_056075.1         --------------EDFKQQ----------VPYKED-----KPQ-----------VPLES      157
NP_001284524.1      --------------EDFKQQ----------VPYKED-----KPQ-----------VPLES      157


NP_001138829.1      QPVEFNHAINYVNKIKNRFQGQPDIYKAFLEILHTYQKEQRNAKEAGGNYTPALTEQEVY      359
NP_001284526.1      ------------------------------------------------------------      0
NP_056075.1         DSVEFNNAISYVNKIKTRFLDHPEIYRSFLEILHTYQKEQLNTR---GRPFRGMSEEEVF      214
NP_001284524.1      DSVEFNNAISYVNKIKTRFLDHPEIYRSFLEILHTYQKEQLNTR---GRPFRGMSEEEVF      214


NP_001138829.1      AQVARLFKNQEDLLSEFGQFLPDANSSVLLSKTTAEKVDSVRNDHGGTVKKPQLNNKPQR      419
NP_001284526.1      ------------------------------------------------------------      0
NP_056075.1         TEVANLFRGQEDLLSEFGQFLPEAKRSLFTGNGPCEMHSVQKNEHDKTPEH---SRKRSR      271
NP_001284524.1      TEVANLFRGQEDLLSEFGQFLPEAKRSLFTGNGPCEMHSVQKNEHDKTPEH---SRKRSR      271


NP_001138829.1      PSQNGCQIRRHPTGTTPPVKKKPKLLNLKDSSMADASKHGGGTESLFFDKVRKALRSAEA      479
NP_001284526.1      ------------------------------------------------------------      0
NP_056075.1         PSL--------LRPVSAPAKKKMKLRGTKDLSIAAVGKYGTLQEFSFFDKVRRVLKSQEV      323
NP_001284524.1      PSL--------LRPVSAPAKKKMKLRGTKDLSIAAVGKYGTLQEFSFFDKVRRVLKSQEV      323


NP_001138829.1      YENFLRCLVIFNQEVISRAELVQLVSPFLGKFPELFNWFKNFLGYKESVHLETYPKERAT      539
NP_001284526.1      ------------------------------------------------------------      0
NP_056075.1         YENFLRCIALFNQELVSGSELLQLVSPFLGKFPELFAQFKSFLGVKELSFA-PPMSDRSG      382
NP_001284524.1      YENFLRCIALFNQELVSGSELLQLVSPFLGKFPELFAQFKSFLGVKELSFA-PPMSDRSG      382


NP_001138829.1      EGIAMEIDYASCKRLGSSYRALPKSYQQPKCTGRTPLCKE--------------------      579
NP_001284526.1      --------------------------------------------------------MQRH      4
NP_056075.1         DGISREIDYASCKRIGSSYRALPKTYQQPKCSGRTAICKELDHWTLLQGSWTDDYCMSKF      442
NP_001284524.1      DGISREIDYASCKRIGSSYRALPKTYQQPKCSGRTAICKE--------------------      422


NP_001138829.1      ------------VLNDTWVSFPSWSEDSTFVSSKKTQYEEHIYRCEDERFELDVVLETNL      627
NP_001284526.1      S----RHFLLVQVLNDTWVSFPSWSEDSTFVSSKKTPYEEQLHRCEDERFELDVVLETNL      60
NP_056075.1         KNTCWIPGYSAGVLNDTWVSFPSWSEDSTFVSSKKTPYEEQLHRCEDERFELDVVLETNL      502
NP_001284524.1      ------------VLNDTWVSFPSWSEDSTFVSSKKTPYEEQLHRCEDERFELDVVLETNL      470
                                ************************ ***:::*****************
```

### Supplementary Figure S2 | Adams et al.

```
NP_001138829.1      ATIRVLEAIQKKLSRLSAEEQAKFRLDNTLGGTSEVIHRKALQRIYADKAADIIDGLRKN    687
NP_001284526.1      ATIRVLESVQKKLSRMAPEDQEKFRLDDSLGGTSEVIQRRAIYRIYGDKAPEIIESLKKN    120
NP_056075.1         ATIRVLESVQKKLSRMAPEDQEKFRLDDSLGGTSEVIQRRAIYRIYGDKAPEIIESLKKN    562
NP_001284524.1      ATIRVLESVQKKLSRMAPEDQEKFRLDDSLGGTSEVIQRRAIYRIYGDKAPEIIESLKKN    530
                    *******::******:: *:* *****::********:*:*: ***.*** :**:.*:**

NP_001138829.1      PSIAVPIVLKRLKMKEEEWREAQRGFNKVWREQNEKYYLKSLDHQGINFKQNDTKVLRSK    747
NP_001284526.1      PVTAVPVVLKRLKAKEEEWREAQQGFNKIWREQYEKAYLKSLDHQAVNFKQNDTKALRSK    180
NP_056075.1         PVTAVPVVLKRLKAKEEEWREAQQGFNKIWREQYEKAYLKSLDHQAVNFKQNDTKALRSK    622
NP_001284524.1      PVTAVPVVLKRLKAKEEEWREAQQGFNKIWREQYEKAYLKSLDHQAVNFKQNDTKALRSK    590
                    *  ***:****** *********:****:**** ** ********.:********.****

NP_001138829.1      SLLNEIESIYDERQEQATEENAGVPVGPHLSLAYEDKQILEDAAALIIHHVKRQTGIQKE    807
NP_001284526.1      SLLNEIESVYDEHQEQHSEGRSAPSSEPHLIFVYEDRQILEDAAALISYYVKRQPAIQKE    240
NP_056075.1         SLLNEIESVYDEHQEQHSEGRSAPSSEPHLIFVYEDRQILEDAAALISYYVKRQPAIQKE    682
NP_001284524.1      SLLNEIESVYDEHQEQHSEGRSAPSSEPHLIFVYEDRQILEDAAALISYYVKRQPAIQKE    650
                    ********:***:*** :* .:.    *** :.***:********** ::**** .****

NP_001138829.1      DKYKIKQIMHHFIPDLLFAQRGDLSDVEEEEEEEMDVDEAT-----GAVKKHNGVGGSPP    862
NP_001284526.1      DQGTIHQLLHQFVPSLFFSQQLDLGASEESADEDRDSPQGQTTDPSERKKPAPGPHSSPP    300
NP_056075.1         DQGTIHQLLHQFVPSLFFSQQLDLGASEESADEDRDSPQGQTTDPSERKKPAPGPHSSPP    742
NP_001284524.1      DQGTIHQLLHQFVPSLFFSQQLDLGASEESADEDRDSPQGQTTDPSERKKPAPGPHSSPP    710
                    *: .*:*::*:*:*.*:*:*: **.  **. :*: *  :.         *   *  .***

NP_001138829.1      KSKLLFSNTAA---------QKLRGMDEVYNLFYVNNNWYIFMRLHQILCLRLLRICSQA    913
NP_001284526.1      EEKGAFGDAPATEQPPLPPPAPHKPLDDVYSLFFANNNWYFFLRLHQTLCSRLLKIYRQA    360
NP_056075.1         EEKGAFGDAPATEQPPLPPPAPHKPLDDVYSLFFANNNWYFFLRLHQTLCSRLLKIYRQA    802
NP_001284524.1      EEKGAFGDAPATEQPPLPPPAPHKPLDDVYSLFFANNNWYFFLRLHQTLCSRLLKIYRQA    770
                    :.*  *.:: *            : :*:**.**:.*****:*:**** ** ***:*  **

NP_001138829.1      ERQIEEENREREWEREVLGIKRDKSDSPAIQLRLKEPMDVDVEDYYPAFLDMVRSLLDGN    973
NP_001284526.1      QKQLLEYRTEKEREKLLCEGRREKGSDPAMELRLKQPSEVELEEYYPAFLDMVRSLLEGS    420
NP_056075.1         QKQLLEYRTEKEREKLLCEGRREKGSDPAMELRLKQPSEVELEEYYPAFLDMVRSLLEGS    862
NP_001284524.1      QKQLLEYRTEKEREKLLCEGRREKGSDPAMELRLKQPSEVELEEYYPAFLDMVRSLLEGS    830
                    ::*: * . *:* *: :   :*:*...**::****:* :*::*:*************:*.

NP_001138829.1      IDSSQYEDSLREMFTIHAYIAFTMDKLIQSIVRQLQHIVSDEICVQVTDLYLAENNNGAT    1033
NP_001284526.1      IDPTQYEDTLREMFTIHAYVGFTMDKLVQNIARQLHHLVSDDVCLKVVELYLNEKKRGAA    480
NP_056075.1         IDPTQYEDTLREMFTIHAYVGFTMDKLVQNIARQLHHLVSDDVCLKVVELYLNEKKRGAA    922
NP_001284524.1      IDPTQYEDTLREMFTIHAYVGFTMDKLVQNIARQLHHLVSDDVCLKVVELYLNEKKRGAA    890
                    ** :****:**********:.******:*.*.***:*:***::*::*.:*** *::.**:

NP_001138829.1      GGQLNTQNSRSLLESTYQRKAEQLMSDENCFKLMFIQSQGQVQLTIELLDTEEENSDDPV    1093
NP_001284526.1      GGNLSSRCVRAARETSYQWKAERCMADENCFKVMFLQRKGQVIMTIELLDTEEAQTEDPV    540
NP_056075.1         GGNLSSRCVRAARETSYQWKAERCMADENCFKVMFLQRKGQVIMTIELLDTEEAQTEDPV    982
NP_001284524.1      GGNLSSRCVRAARETSYQWKAERCMADENCFKVMFLQRKGQVIMTIELLDTEEAQTEDPV    950
                    **:*.::  *:  *::** ***: *:******:**:* :*** :********* :::***

NP_001138829.1      EAERWSDYVERYMNSDTTSPELREHLAQKPVFLPRNLRRIRKCQRGREQQEKEGKEGNSK    1153
NP_001284526.1      EVQHLARYVEQYVGTEGASSSPTEGFLLKPVFLQRNLKKFRRRWQSEQARALRGEARSSW    600
NP_056075.1         EVQHLARYVEQYVGTEGASSSPTEGFLLKPVFLQRNLKKFRRRWQSEQARALRGEARSSW    1042
NP_001284524.1      EVQHLARYVEQYVGTEGASSSPTEGFLLKPVFLQRNLKKFRRRWQSEQARALRGEARSSW    1010
                    *.:: : ***:*:.:: :* .  * :  ***** ***:::*:  :..: :  .*:  .*

NP_001138829.1      KTMENVDSLDKLECRFKLNSYKMVYVIKSEDYMYRRTALLRAHQSHERVSKRLHQRFQAW    1213
NP_001284526.1      KRLVGVESACDVDCRFKLSTHKMVFIVNSEDYMYRRGTLCRAKQVQPLVLLRHHQHFEEW    660
NP_056075.1         KRLVGVESACDVDCRFKLSTHKMVFIVNSEDYMYRRGTLCRAKQVQPLVLLRHHQHFEEW    1102
NP_001284524.1      KRLVGVESACDVDCRFKLSTHKMVFIVNSEDYMYRRGTLCRAKQVQPLVLLRHHQHFEEW    1070
                    * : .*:*  .::*****.::***::::******** :* **:* :  *  * **:*: *

NP_001138829.1      VDKWTKEHVPREMAAETSKWLMGEGLEGLVPCTTTCDTETLHFVSINKYRVKYGTVFKAP    1273
NP_001284526.1      HSRWLEDNVTVEAASLVQDWLMGEEDEDMVPCKTLCETVHVHGLPVTRYRVQYSRRPASP    720
NP_056075.1         HSRWLEDNVTVEAASLVQDWLMGEEDEDMVPCKTLCETVHVHGLPVTRYRVQYSRRPASP    1162
NP_001284524.1      HSRWLEDNVTVEAASLVQDWLMGEEDEDMVPCKTLCETVHVHGLPVTRYRVQYSRRPASP    1130
                     .:* :::*  * *: ...*****  *.:***.* *:*  :* : :.:***:*.    :*
```

### Supplementary Figure 3 | Adams et al.

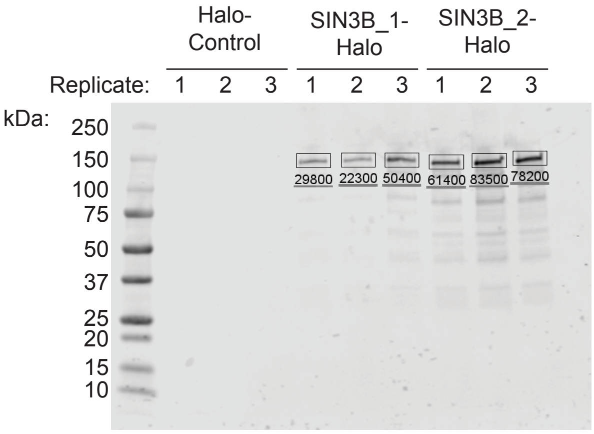

### Supplementary Figure S4 | Adams et al.

**Needle Pairwise Sequence Alignment**

```
Protein           Accession
RBBP4 isoform a: NP_005601.1
RBBP4 isoform c: NP_001128728.1


Program: needle
        -datafile EBLOSUM62
        -gapopen 10.0
        -gapextend 0.5
        -endopen 10.0
        -endextend 0.5
        -Matrix: EBLOSUM62
        -Gap_penalty: 10.0
        -Extend_penalty: 0.5

Length: 425
Identity:     390/425 (91.8%)
Similarity:   390/425 (91.8%)
Gaps:          35/425 ( 8.2%)
Score: 2113.0


NP_005601.1        1 MADKEAAFDDAVEERVINEEYKIWKKNTPFLYDLVMTHALEWPSLTAQWL     50
                                                       |||||||||||||||
NP_001128728.      1 -----------------------------------MTHALEWPSLTAQWL     15

NP_005601.1       51 PDVTRPEGKDFSIHRLVLGTHTSDEQNHLVIASVQLPNDDAQFDASHYDS    100
                     ||||||||||||||||||||||||||||||||||||||||||||||||||
NP_001128728.     16 PDVTRPEGKDFSIHRLVLGTHTSDEQNHLVIASVQLPNDDAQFDASHYDS     65

NP_005601.1      101 EKGEFGGFGSVSGKIEIEIKINHEGEVNRARYMPQNPCIIATKTPSSDVL    150
                     ||||||||||||||||||||||||||||||||||||||||||||||||||
NP_001128728.     66 EKGEFGGFGSVSGKIEIEIKINHEGEVNRARYMPQNPCIIATKTPSSDVL    115

NP_005601.1      151 VFDYTKHPSKPDPSGECNPDLRLRGHQKEGYGLSWNPNLSGHLLSASDDH    200
                     ||||||||||||||||||||||||||||||||||||||||||||||||||
NP_001128728.    116 VFDYTKHPSKPDPSGECNPDLRLRGHQKEGYGLSWNPNLSGHLLSASDDH    165

NP_005601.1      201 TICLWDISAVPKEGKVVDAKTIFTGHTAVVEDVSWHLLHESLFGSVADDQ    250
                     ||||||||||||||||||||||||||||||||||||||||||||||||||
NP_001128728.    166 TICLWDISAVPKEGKVVDAKTIFTGHTAVVEDVSWHLLHESLFGSVADDQ    215

NP_005601.1      251 KLMIWDTRSNNTSKPSHSVDAHTAEVNCLSFNPYSEFILATGSADKTVAL    300
                     ||||||||||||||||||||||||||||||||||||||||||||||||||
NP_001128728.    216 KLMIWDTRSNNTSKPSHSVDAHTAEVNCLSFNPYSEFILATGSADKTVAL    265

NP_005601.1      301 WDLRNLKLKLHSFESHKDEIFQVQWSPHNETILASSGTDRRLNVWDLSKI    350
                     ||||||||||||||||||||||||||||||||||||||||||||||||||
NP_001128728.    266 WDLRNLKLKLHSFESHKDEIFQVQWSPHNETILASSGTDRRLNVWDLSKI    315

NP_005601.1      351 GEEQSPEDAEDGPPELLFIHGGHTAKISDFSWNPNEPWVICSVSEDNIMQ    400
                     ||||||||||||||||||||||||||||||||||||||||||||||||||
NP_001128728.    316 GEEQSPEDAEDGPPELLFIHGGHTAKISDFSWNPNEPWVICSVSEDNIMQ    365

NP_005601.1      401 VWQMAENIYNDEDPEGSVDPEGQGS    425
                     |||||||||||||||||||||||||
NP_001128728.    366 VWQMAENIYNDEDPEGSVDPEGQGS    390


#---------------------------------------
#---------------------------------------
```

### Supplementary Figure S5 | Adams et al.

(A)

```
         10        20        30        40        50        60
....|....|....|....|....|....|....|....|....|....|....|....|
ATGCATCCTGCAAGCGCATAGGATCCAGCTACCGGGCACTCCCCAAAACCTACCAGCAGC 60

         70        80        90       100       110       120
....|....|....|....|....|....|....|....|....|....|....|....|
CCAAGTGCAGTGGGAGGACAGCCATCTGCAAGGAGCTTGACCATTGGACACTTCTCCAGG 120

        130       140       150       160       170       180
....|....|....|....|....|....|....|....|....|....|....|....|
GTTCGTGGACAGACGATTACTGCATGTCCAAGTTCAAGAATACCTGCTGGATTCCAGGAT 180

        190       200       210       220       230       240
....|....|....|....|....|....|....|....|....|....|....|....|
ATAGTGCAGGGGTACTGAACGACACCTGGGTCTCCTTCCCTTCCTGGTCTGAGGACTCCA 240

        250       260       270
....|....|....|....|....|....|
CGTTCGTCAGCTCCAAGAAGACACCGTACG 270
```

(B)

```
         10        20        30        40        50        60
....|....|....|....|....|....|....|....|....|....|....|....|
AAAAGCGATCGCCATGCAGCGTCATTCACGGCATTTCCTCTTGGTGCAGGTACTGAACGA 60

         70        80        90       100       110       120
....|....|....|....|....|....|....|....|....|....|....|....|
CACCTGGGTCTCCTTCCCTTCCTGGTCTGAGGACTCCACGTTCGTCAGCTCCAAGAAGAC 120

        130
....|....|..
ACCGTACGAAAA 132
```
